# Selective Optogenetic Stimulation of Glutamatergic, but not GABAergic, Vestibular Nuclei Neurons Induces Immediate and Reversible Postural Imbalance in Mice

**DOI:** 10.1101/2020.09.03.281980

**Authors:** Q. Montardy, M. Wei, T. Yi, X. Liu, Z. Zhou, J. Lai, S. Besnard, B. Tighilet, C. Chabbert, L. Wang

## Abstract

Glutamatergic and GABAergic neurons represent the neural components of the medial vestibular nuclei. We assessed the functional role of glutamatergic and GABAergic neuronal pathways arising from the vestibular nuclei (VN) in the maintenance of gait and balance by optogenetically stimulating the VN in VGluT2-cre and GAD2-cre mice. We demonstrate that glutamatergic, but not GABAergic VN neuronal subpopulation is responsible for immediate and strong posturo-locomotor deficits, comparable to unilateral vestibular deafferentation models. During optogenetic stimulation, the support surface dramatically increased in VN^VGluT2+^ mice, and rapidly fell back to baseline after stimulation, whilst it remained unchanged during similar stimulation of VN^GAD2+^ mice. This effect persisted when vestibular compensation was removed. Posturo-locomotor alterations evoked in VN^VGluT2+^ animals were still present immediately after stimulation, while they disappeared 1h later. Overall, these results indicate a fundamental role for VN^VGluT2+^ neurons in balance and posturo-locomotor functions, but not for VN^GAD2+^ neurons, in this specific context. This new optogenetic approach will be useful to characterize the role of the different VN neuronal populations involved in vestibular physiology and pathophysiology.

**Highlights:**

- For the first time, Vestibular nuclei were optogenetically stimulated in free-moving animals, to asses for glutamatergic and GABAergic neurons functions in posturo-locomotor behaviors.
- Brief optogenetic activation of VNVGluT2+, but not VNGAD2+, induced immediate and strong postural deficit.
- Stimulation of VNVGluT2+ neurons provoked an imbalance with continuous effect on locomotion for a short period of time after stimulation.
- These results are comparable to the classical vestibular deafferentation models during their peak of deficit, and set optogenetic stimulation as a new model to study vestibular deficits.

## Introduction

The first mentions of vertigo as a specific symptom date back to antiquity with Hippocrates writings (Adams, 1849), and it was not until the beginning of the 19^th^ century that the study of vertigo and balance entered the experimental phases, with the works of Purkinje in humans (Purkinje, 1819), and Flourens in the pigeon (Flourens, 1842). These led to the establishment of a direct link between damage to the inner-ear sensors and the characteristic posturo-locomotor and vestibulo-ocular deficits encountered in dizzy patients. A century later, Lorente de No introduced the first anatomical descriptions of the central pathways that connect the peripheral vestibular sensors to the oculomotor muscles (Lorente De No, 1933).

The anatomical and functional organization of the vestibular system is now better understood, and its contribution to a wide range of functions, from postural and oculomotor reflexes to spatial representation and cognition, has been underscored (Angelaki and Cullen, 2008; Borel et al., 2008). Head accelerations resulting from the interaction with our environment are detected by the mechanosensitive hair cells located in the inner ear sensory organs, and encoded into an electrical signal. This sensory information is then conveyed to the brain stem vestibular nuclei (VN) through the vestibular nerve. Four vestibular nuclei (VN) are located in the brainstem under the floor of the IV^th^ ventricle: the median (MVN), inferior VN (IVN), lateral (LVN) and superior (SVN) vestibular nuclei. At the VN level, this essentially sensory information is integrated and transformed into premotor signals, the transport of which is ensured by vestibulo-spinal and vestibulo-oculomotor pathways. VN Neurons project, through the medial longitudinal fasciculus (MLF), onto the oculomotor nuclei whose motoneurons control gaze stabilization during head movements. The vestibular nuclei influence postural control via two descending pathways to the spinal cord, the lateral (mostly from ipsilateral NVL neurons) and medial (mostly from ipsi and contralateral NVM neurons) vestibulospinal tracts (Wilson and Jones, 1979). Vestibular information is also transferred via vestibulo-thalamo-cortical pathways to different cortical areas providing perceptual and cognitive functions such as perception and orientation of body in space (Lopez, 2016; Lopez et al., 2015). Finally, the vestibular nuclei are interconnected to a multitude of neurovegetative centers in the brainstem (Balaban and Beryozkin, 1994; Balaban and Porter, 1998). Given this anatomo-functional organization of the vestibular system and its projection targets, unilateral alteration of peripheral vestibular inputs induces a quadruple syndrome: posturo-locomotor, oculomotor, vegetative and perceptivo-cognitive. Symptoms progressively improve, each with its own kinetics, generally leading to an almost full disappearance of the syndrome. This behavioral recovery phenomenon is referred to as “vestibular compensation” (Lacour et al., 2016).

### Animal models in vestibular pathologies

Different animal models of vestibular disorders have been developed. Researchers set out to determine either the pathogenic conditions likely to induce vestibular dysfunction and its functional consequences on the basis of an epidemiological rationale, or the neurophysiological mechanisms likely involved. The first case included: 1) ototoxic-based vestibular damage, reproducing inner ear-specific toxicity of drugs such as aminoglycosides or cisplatin, and food-borne ototoxicity (Hirvonen et al., 2005; Vignaux et al., 2012; Xia et al., 2012; Zhang et al., 2003); 2) excitotoxically-induced vestibular damage, reproducing the deleterious consequences on the first synapse of sensory cells suffering (Brugeaud et al., 2007; Desmadryl et al., 2012; Dyhrfjeld-Johnsen et al., 2013; Gaboyard-Niay et al., 2016; O’Neill et al., 1999); 3) destruction of peripheral receptors through surgical labyrintectomy (Hitier et al., 2010; Patk et al., 2003; Smith and Curthoys, 1989); 4) vestibular neurectomy that relates to the full section of the vestibular nerve. This later operation can be achieved either peripheral unilateral vestibular neurectomy (peripheral UVN; nerve section between vestibular endorgans and Scarpa’s ganglion) (Hamann et al., 1998; Li et al., 1995) or central UVN (between the Scarpa’s ganglion and the brainstem). Peripheral unilateral vestibular neurectomy (UVN) leads to the loss of inputs from vestibular endorgans, whilst preserving those arising from vestibular ganglion neurons. Central UVN induces complete loss of inputs arising from both vestibular endorgans and Scarpa ganglion neurons. Several examples of central UVN have been documented previously in different species such as monkeys (Lacour et al., 1976), cats (Xerri and Lacour, 1980) and rats (Li et al., 1995). Along with the lesion models mentioned above, there are models of reversible modulation of the vestibular input. The first to be developed was the so-called “vestibular anesthesia” (Bárány, 1936). Assuming that the episodes of vertigo attacks observed in Menière patients resulted from transient unilateral hyperexcitability of primary vestibular neurons, this approach consisted of counteracting this hyperexcitability through transtympanic administration of neuronal activity, such as lidocaine. A similar modulatory effect was obtained through transtympanic administration of tetrodotoxin (TTX), a selective blocker of voltage-gated calcium channels (Dutheil et al., 2011). Under these conditions, pharmacological modulation of peripheral sensory input either reproduced in animals the vestibular syndrome encountered in patients with acute peripheral vestibulopathy, or led to alleviation of symptoms when administrated to the ear opposite to the damaged one.

Although these models have led to a better understand of the conditions that generate vertigo syndrome, and its temporal evolution, questions related to the anatomical and functional organization of the vestibular central pathways remain. As an example, the target cerebral structures of neurons projecting from the VN have are not all been identified, in particular in the cortex. Likewise, the central pathways projecting from cortical areas towards the VN are poorly documented. This lack of data prevents the development of studies attempting to understand the links between vestibular, cognitive and emotional functions. To answer these questions, we used optogenetics in mice to stimulate glutamatergic and GABAergic VN neuronal subpopulations. The principle of optogenetics is based on the insertion of genes encoding photoactive proteins, such as opsins, into defined cell populations (VGluT2+ and GAD2+). This method makes it possible to specifically stimulate these neurons. Our results demonstrate that it is possible to use optogenetics to generate posturo-locomotor deficits, an essential component of the vestibular syndrome. Here, these posturo-locomotor deficits are compared with those induced by classical chemical labyrinthectomy.

## Results

### Validation of the optogenetic method in vestibular nuclei

Mice expressing either a cre recombinase driven by VGluT2+ (vesicular glutamate transporter 2) promoter in glutamatergic neurons (VGluT2-ires-cre), or a cre recombinase driven by GAD2 (glutamate decarboxylase 2) promoter in GABAergic neurons (GAD2-ires-Cre) were injected uniterally with the cre-dependent virus AAV9-DIO-ChR2-mCherry in the vestibular nuclei (VN). In parallel, a control group of mice expressing VGluT2-cre or GAD2-cre received injections of AAV9-DIO-mCherry at the same location. Three weeks after injection, all mice were implanted with an optical fiber above the VN for the purpose of vestibular optical stimulation on awake and free-moving animals (Fig.1A,B). After completion of all experimental tests, brains were extracted and sectioned to verify viral expression and location of optical fibers (VGluT2-cre: Fig.1C,D; GAD2-cre: Fig.1E,F) and data was only used for animals when both parameters were verified. With regard to viral expression, VN neurons that co-expressed mCherry viral fluorescence and DAPI staining indicated that VN neurons were locally infected in both VGluT2-cre (Fig.1C, right) and GAD2-cre (Fig.1E, right) groups. Optical fibers were implanted in the upper parvicellular part of medial VN (Fig.1 C,E,D,F).

**Figure 1.**
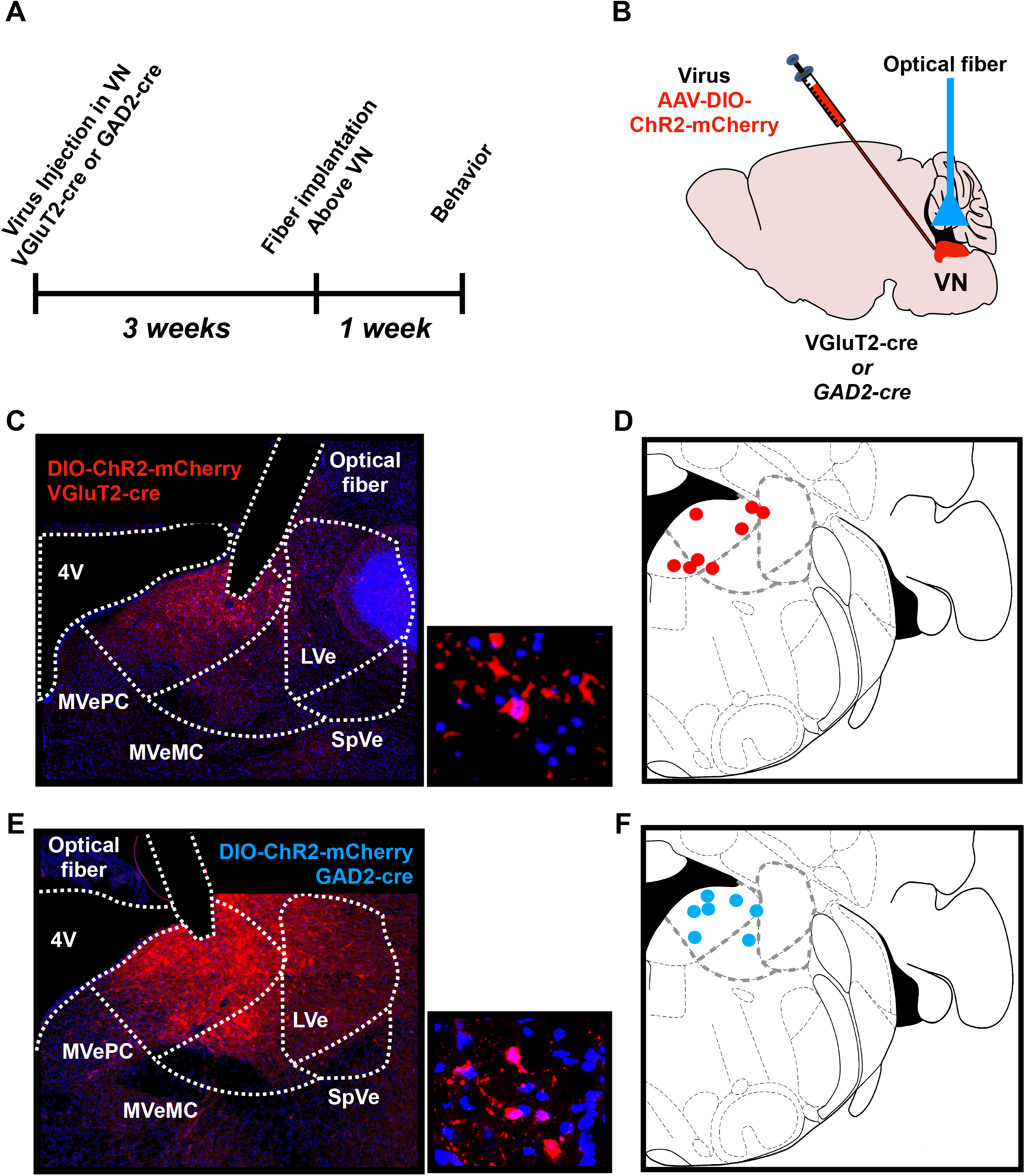
VN virus injection and optical fiber implantation. **A.** Schematic showing the experimental protocol. AAV9-DIO-ChR2-mCherry was injected int the VN of VGluT2-ires-cre or GAD2-ires-cre. Three weeks later, an optical fiber was positioned in the vicinity of the VN, and behavioral tests were performed 1 week later. **B.** Schematic representation of virus injection and optical fiber location in the VN. **C.** Representative photograph of virus expression in the VN of VGluT2+ mice. Red, ChR2-mCherry virus; Blue, DAPI. Scale bar, 100 μm (left), and 50 μm (right). MVePC (parvicellular medial vestibular nucleus) MVeMC (magnoellular medial vestibular nucleus), SPVe (superior vestibular nucleus), LVe (lateral vestibular nucleus) **D.** Representation of fiber tip position for individual VGluT2+ animals (in red) **E.** Representative picture of virus expression in VN of GAD2+ animals. **F.** Representation of fiber tip position for individual VGluT2+ animals (in blue).

### Unilateral optogenetic stimulation of VN^VGluT2+^ neurons, but not GAD2 neurons, triggers strong posturo-locomotor deficits

Before starting the optogenetic stimulation protocol, one group underwent a unilateral vestibular deafferentation using chemical labyrintectomy in order to compare the functional consequences of optogenetic stimulation with characteristic acute vestibular syndrome. This was done by transtympanic administration of sodium arsanilate in WT mice, as previously reported in rats (Vigneaux et al. 2012), and posturo-locomotor behavior was recorded and analyzed in an open field using both subjective (Pericat et al. 2017) and automatized (Rastoldo et al. 2020) analyses. Posturo-locomotor behavioral parameters, included head tilt, bobbing, circling, retropulsion and tumbling (Fig. 2A) were observed at several time points over 3 weeks (Fig. 2B, gray). All mice (n=6) exhibited specific vestibular deficits, such as head tilt and circling, both oriented towards the injured side. In addition, mice exhibited limb extension towards the intact side, accompanied by limb flexion on the opposite side, often leading to falls towards the lesion side. Animals were strongly and significantly affected on the first few days post-lesion, especially at Day 2 (Baseline = 0 vs. D2 = 8.17, P<0.05, t=2.69, df=8, n=6), then partially recovered from D3 and over the 3 weeks (D2 vs. W3, P<0.05, t=2.94, df=5, Fig. 2.B). A relative heterogeneity of deficits between animals emerged following a qualitative analysis of D2 data, with some showing only head tilt, whilst others displaying all categories of symptoms, including tumbling (Fig. 2.C).

**Figure 2.**
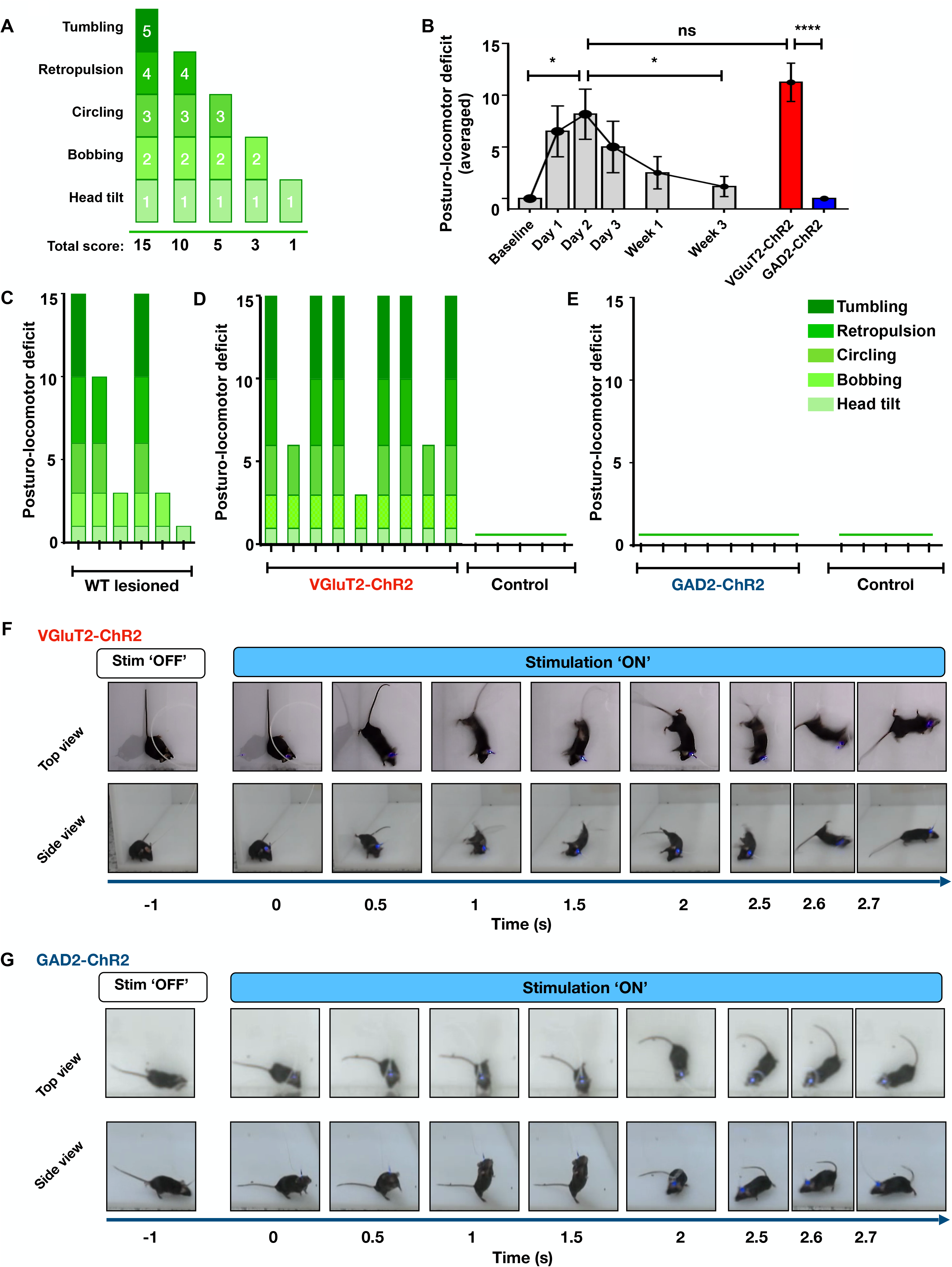
Optogenetic stimulation of VN^VGluT2+^ neurons, but not GAD2 neurons, induced strong posturo-locomotor deficits. **A.** Evaluation grid used to score behavior in the open field. If all symptoms are expressed, then a rating of 15 is scored, if all except tumbling are expressed, the rating 10 is scored, and so on. **B.** Illustration of the time course of the behavioral evaluation score for WT animals with arsanilate lesions (gray), compared with VN^VGluT2+^ (red) and VN^GAD2+^ mice during optogenetic stimulation. **C.** Behavioral evaluation score of WT-lesion animals, at D2 post-lesion, whilst animals were placed on a static ground. **D.** Behavioral evaluation score for the VGluT2-ChR2 (left) and VGluT2-mCherry (right) groups. **E.** GAD-ChR2 (left) and GAD2-mCherry (right) groups during a 30 s stimulation whilst animals were on the ground. GAD-ChR2 and mCherry do not show a deficit. **F.** Example of posturo-locomotor behavior from a VGluT2-ChR2 mouse (essentially tumbling) **G.** Illustration of a GAD-ChR2 mouse during stimulation.

Mice in the VGluT2-cre ChR2 and control mCherry groups were placed in an open field and observed for 5 minutes during which animals did not show any sign of posturo-locomotor deficit. During persistent optogenetic stimulation (20 Hz, 5 ms, 473 nm for 30 s) that followed, the VGluT2-ChR2 group displayed strong posturo-locomotor deficits similar to that observed in the arsanilate group (Fig. 2D, left). However, unlike mice in the arsanilate group, all VGluT2-ChR2 mice exhibited limb extension on the stimulated side, accompanied by limb flexion on the non-stimulated side. Six of nine mice exhibited instantaneous tumbling towards the non-stimulated side (Fig. 2D, 2F, Sup. Video 1). In addition, all mice exhibited circling behavior oriented towards the stimulated side. In comparison, VGluT2-mCherry control group animals did not show deficits during the entire stimulation period (Fig. 2D, right). Quantitatively, the VGluT2-ChR2 group tended to present a higher deficit than the arsanilate group, but not significantly different to D2 post-lesion arsanilate animals (D2 = 8.17 vs. VGluT2-ChR2 = 11.67, P>0.1; Fig. 2B, red). We then followed the same procedure using GAD2-cre animals and stimulated VN GABAergic neurons (60 Hz, 5 ms, 473 nm for 30 s) and did not find any postural deficits in the ChR2 or mCherry animals (Fig. 2E). This is as also demonstrated by video examples (Fig. 2G). In general, the postural behavior in the GAD2-ChR2 mice was comparable to the non-lesion mice (GAD2-ChR2 = 0 vs. VGluT2-ChR2 = 11.67, P<0.001, t=6.48, df=15; Fig. 2B, blue). These data indicate that activation of VN glutamatergic neurons leads to immediate and strong posturo-locomotor deficits that are, on average, comparable to those observed for arsanilate group at day 2. In contrast, activation of GABAergic neurons does not appear to have such a marked effect.

### Tail-hanging landing tests confirmed that the VN^VGluT2+^ population is responsible for posturo-locomotor deficits, compared with the VN^GAD2+^ population

Above, we show that stimulation of VN^VGluT2+^ (but not VN^GAD2+^) neurons led to posturo-locomotor deficits whilst the animals were standing on a static ground. To verify whether this observation persists when vestibular compensation is removed, we performed a tail-hanging landing test and scored three mains parameters: landing, twirl, and posture after landing (Fig. 3A). First, WT animals following arsanilate injection showed strong and homogeneous deficits. Compared to baseline measures, twirls and posture landing were affected (mean score from all mice=6), whilst posture after landing was only moderately affected (mean score for 4/6 mice=1; Fig. 3C). Across testing, effects were especially marked at D2 post-lesion (Baseline = 0 vs. D2 = 7.33, P<0.0001, t=27.8, df=8, n=6) and did not recover even 3 weeks after surgery (D2 vs. W3, P<0.005, t=4.83, df=5, Fig 3B, gray). Next, mice in the VN^VGluT2+^ and VN^GAD2+^ groups were tested similarly to the WT/arsanilate group and were given 30 s continuous optogenetic stimulation. During stimulation, all VGlut2-ChR2 mice showed the full range of postural deficits, some stronger even than those observed in the WT/arsanilate group, confirming the posturo-locomotor results above, with the exception of one animal that only showed moderate symptoms and no deficit during landing (Fig. 3D, left, Fig. 3F). Control VGluT2-mCherry animals did not show any deficits during light stimulation (Fig. 3D, right), confirming that viral injection surgery did not damage the VN. Overall, stimulation of VN^VGluT2+^ resulted in postural deficits similar to those with arsanilate animals at D2 post-lesion (D2 = 7.33 vs. W3 = 7.78, P>0.1; Fig. 3B, red). Next, GAD2-ChR2 animals were tested and no variation of postural behavior was observed following light stimulation; the GAD2-ChR2 group remained indistinguishable from control GAD2-mCherry group (Fig. 3E, G). Stimulation of GAD2-ChR2 was, overall, similar to arsanilate animals before lesion. Together, these data indicate that brief activation of VN^VGluT2+^ induced strong postural imbalance, whilst transient activation of VN^GAD2+^ neurons do not appear to participate in balance, at least in the short term.

**Figure 3.**
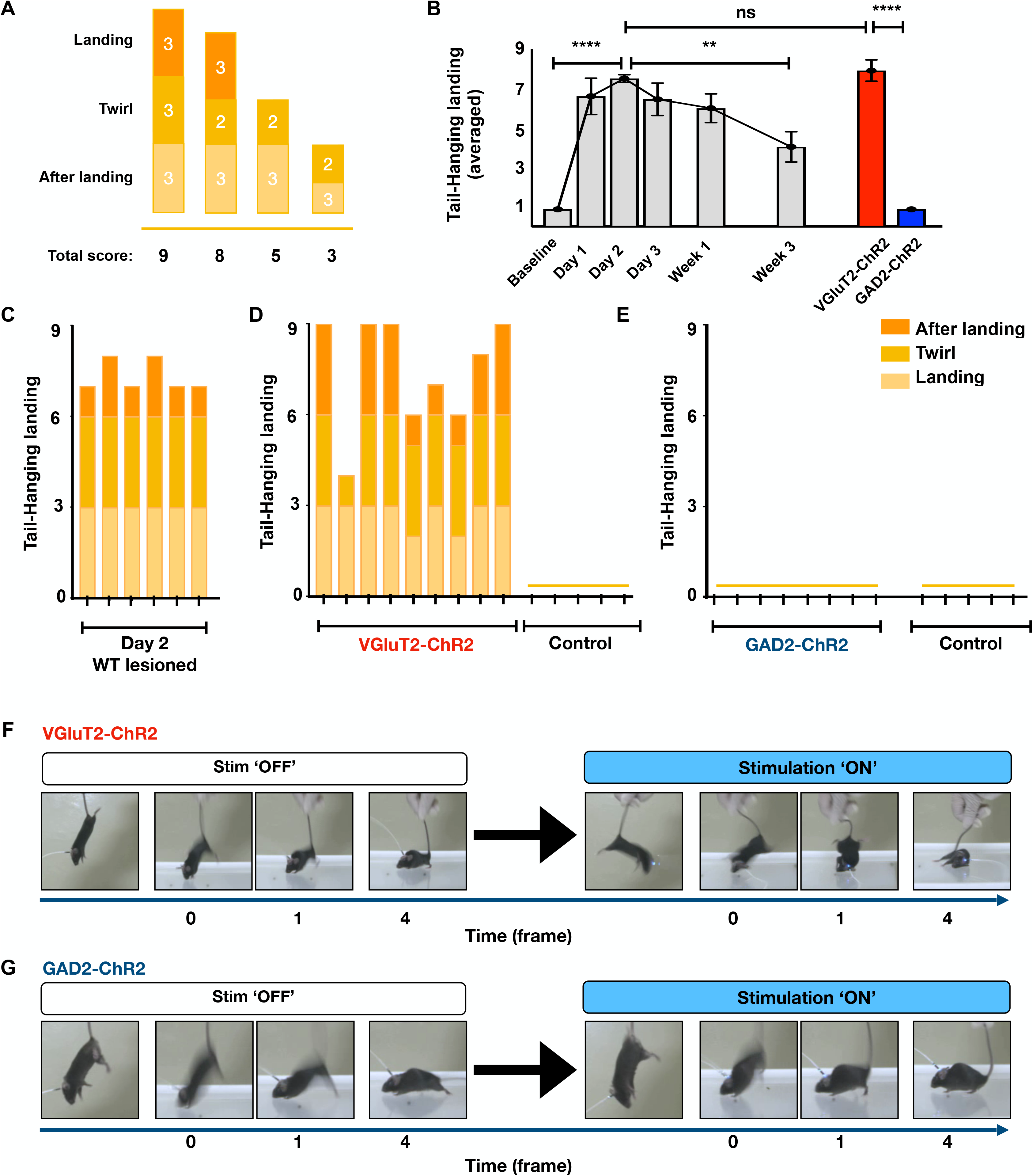
Tail-hanging landing test confirms that the VN^VGluT2+^ population is mainly responsible of posturo-locomotor deficits, compared to the VN^GAD2+^ population. **A.** Evaluation grid used for tail-hanging landing tests. **B.** Time course of the behavioral evaluation score for WT animals with arsanilate lesions (gray), compared to VN^VGluT2+^ (red) and VN^GAD2+^ mice during optogenetic stimulation. **C.** Behavioral evaluation score of the WT-lesion group at D2 post-lesion, whilst undergoing tail-hanging landing tests. **D.** Behavioral evaluation scores of the VGluT2-ChR2 (left), VGluT2-mCherry (right) groups. **E.** GAD-ChR2 (left) and GAD2-mCherry (right) groups during a 30 s stimulation during tail-hanging landing test. The GAD-ChR2 and mCherry groups did not show a deficit. **F.** Example of posturo-locomotor behavior in a VGluT2-ChR2 mouse. **G.** Representative GAD-ChR2 mouse during stimulation. Time in frame: left, before stimulation; right, during stimulation.

### Optogenetic stimulation of VN^VGluT2+^ induced reversible postural instability

For a more thorough assessment of static postural instability caused by VN stimulation, we monitored the support surface area (Marouane et al., 2020) before, during and 5 minutes after optogenetic stimulation. The support surface area was calculated by measuring the surface delimited by the four legs of each mouse whilst they were static for at least one second. Before stimulation or lesion, VGluT2-ChR2, GAD2-ChR2, and WT-lesion animals had similar scores (Fig. 4A, Pre). During optogenetic stimulation, VGluT2+ mice drastically increased their support surface area compared to pre-stimulation (VGluT2-Pre=0.006% vs. VGluT2-Stim=0.0145%, P<0.0001, t=13.66, df=7), and compared to WT-lesion at D2 (VGluT2-Stim=0.0145% vs. D2=0.0113%, P<0.05, t=2.124, df=12; Fig. 4 A, B, D). In line with previous results, GAD2-ChR2 stimulation did not cause any significant difference in support surface area, and was significantly lower than the support surface area of VGluT2-ChR2 during stimulation (VGluT2-Stim=0.0145% vs. GAD2=0.0055%, P<0.0001, t=10.76, df=14, Fig. 4A, Stim; Fig. 4C). At five minutes post-stimulation, higher support surface area scores were still observed in the VGluT2-ChR2 group than in the GAD2-ChR2 group (VGluT2-Stim=0.0068% vs. GAD2=0.0054 %, P<0.05, t=2.49, df=14; Fig. 4A, C. However, VGluT2-ChR2 group scores post-stimulation were not significantly different from pre-stimulation baseline scores. These results confirm that stimulation of VN^VGluT2+^, but not VN^GAD2+^, artificially induced continuous instability during the entire stimulation duration, comparable to and even stronger than that observed in the unilateral vestibular deafferentation model. Importantly, this effect was reversible and faded some minutes after stimulation ended.

**Figure 4.**
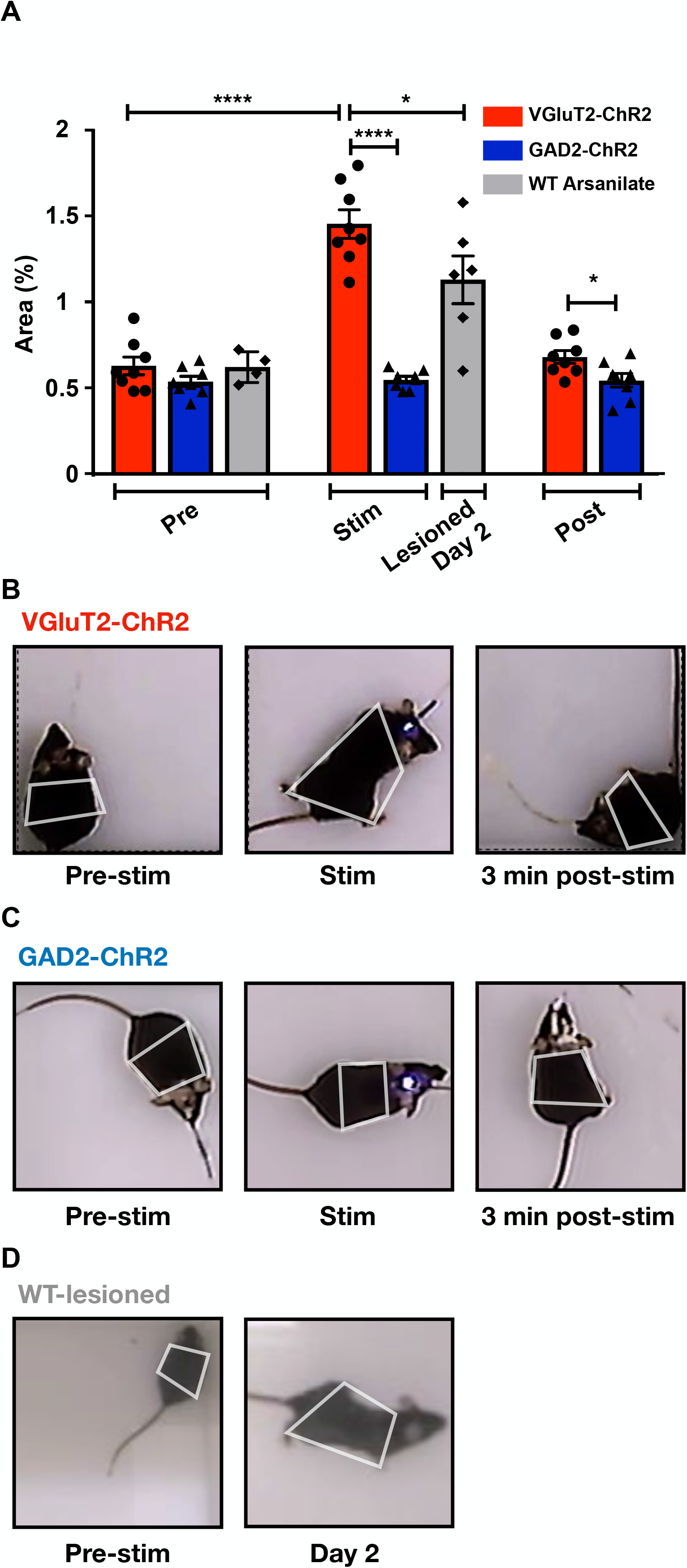
Reversible postural instability upon optogenetic stimulation of VN^VGluT2+^ neurons. **A.** Support surface area as a percentage, averaged across all VN^VGluT2+^ (red), VN^GAD2+^ (blue) and WT-lesion animals (gray). Three 5-min time points were investigated: before stimulation, 5 min after stimulation, and 1 h after stimulation. **B.** Example of the support surface area computed on one VN^VGluT2+^ mouse and **C.** a VN^GAD2+^ mouse, before, right after, and 1 h after stimulation, as well as **D.** WT-lesion animal before and after lesion.

### VN^VGluT2+^ activation induces short-term disturbances of locomotor function

To assess the functional consequences of unilateral optogenetic stimulation of VN on locomotor function, we next studied several parameters previously identified as biomarkers of rodent locomotor dysfunction (Rastoldo et al., 2020). These parameters were monitored in mice placed in an open field before, 5 minutes after, and 1 hour after optogenetic stimulation. Before stimulation, VGluT2-ChR2 and VGluT2-mCherry animals traveled a similar distance compared to immediately after stimulation. However, at 5 minutes post-stimulation, the distance traveled by the VGluT2-ChR2 group was significantly lower than that traveled by the VGluT2-mCherry control group (ChR2=2.04 vs. mCherry=5.87, P<0.001, t=4.7, df=11, Fig. 5A). After 1 hour, VGluT2-ChR2 displacement was not significantly different to the VGluT2-mCherry group. In parallel, the velocity of the VGluT2-ChR2 right after stimulation was significantly lower than the control VGluT2-mCherry group (ChR2=0.012 vs. mCherry=0.019, P<0.001, t=4.6, df=11). This difference disappeared 1 hour after stimulation, where again, the VGluT2-ChR2 group tended to have lower velocity (Fig. 5B). When observing freezing time, the VGluT2-ChR2 group spent significantly more time immobile than the control VGluT2-mCherry group following optogenetic stimulation (ChR2=86.46% vs. mCherry=53.15%, P<0.005, t=3.69, df=11), however, 1 hour later the two groups were not significantly different but there was a trend for higher freezing in the VGluT2-ChR2 group (Fig. 5C). Overall, these results indicate that stimulation of VN^VGluT2+^ significantly altered locomotion up to 5 minutes after stimulation, but this effect disappeared after 1 hour. In comparison, there was no significant difference or trend between GAD2-ChR2 and GAD2-mCherry groups, either for the distanced traveled (Fig. 5D), mean velocity (Fig. 5E) or immobility time (Fig. 5F) at any time point of the experiment. Together, these results corroborate our previous experiment, and indicate that stimulation of VN^VGluT2+^ neurons, but not VN^GAD2+^, continued to have an effect on locomotion for a short period of time after stimulation.

**Figure 5.**
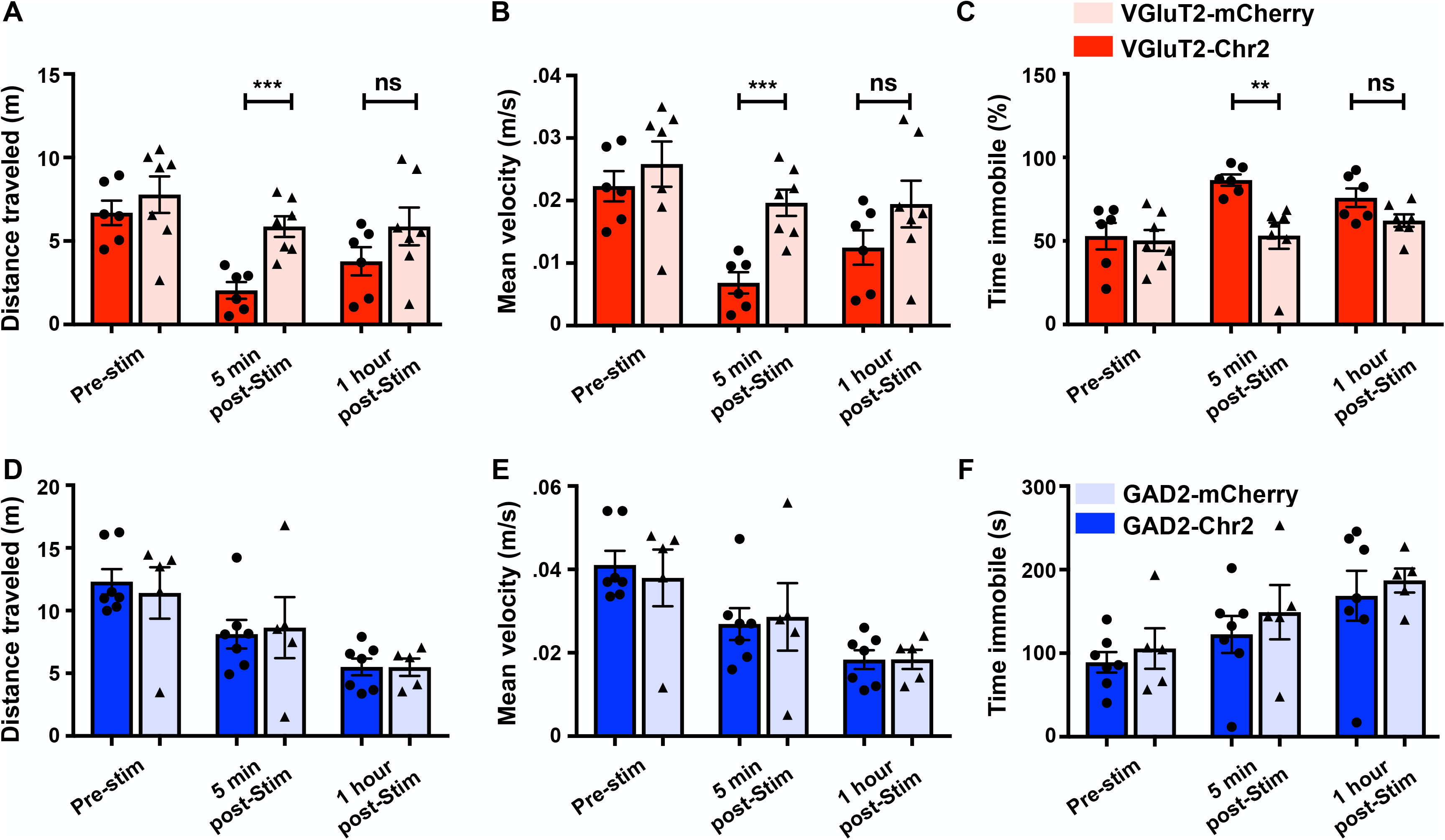
VN^VGluT2+^ activation induces short-term disturbances of locomotor function. Averaged locomotor behaviors during free movement in an open field of mice that received either VGluT2-ChR2 (dark red) or VGluT2-mCherry injection (light red) in the VN. Three 5-min time points were investigated: before stimulation, 5 min after stimulation, and 1 h after stimulation. **A.** Distance moved in meters. **B.** Mean velocity during movement in m/s. **C.** Time spent immobile whilst in the apparatus expressed as a percentage of total experiment time. Averaged locomotor behaviors for GAD2-ChR2 (dark blue) or GAD2-mCherry injection (light blue), computed as previously described **D.** Distance moved. **E.** Mean velocity. **F.** Percentage of time spent immobile.

## Discussion

This study demonstrates for the first time that it is possible to induce reversible posturo-locomotor deficits characteristic of vestibular syndrome by optogenetic stimulation of the vestibular nuclei (VN) area in rodents. Optogenetic stimulation of the glutamatergic VN, but not GABAergic, neuronal subpopulation was responsible for immediate and strong posturo-locomotor deficits, comparable to our unilateral vestibular deafferentation model. Optogenetic stimulation reproduced the mirror image of typical posturo-locomotor behavior evoked by unilateral lesion of the vestibular system. By selective and reproducible stimulation of the different neuronal populations in the vestibular nuclei, this approach brings new opportunities to identify downstream pathways of VN neurons, and to better understand their functional relevance under normal or pathological conditions.

### Technical considerations

Among the four main vestibular nuclei, we chose to stimulate specifically the medial vestibular nuclei (MVN) for various reasons. The medial vestibular nucleus is the one most extensively studied in animal anatomo-functional studies (for review see: Paterson et al., 2004). It is the largest nuclei form the vestibular nuclear complex (VNC) in most mammals (Alvarez et al., 1998) and is known to receive convergent semicircular and otolithic afferents, and to be involved in both oculomotor and postural functions (Wilson and Jones, 1979). The MVN contains a wide diversity of neuronal classes, which project to the oculomotor nuclei, spinal cord, cerebellum, thalamus, contralateral vestibular nuclei, or function as interneurons (Highstein and Holstein, 2006; Straka et al., 2005). It contains both glutamatergic and GABAergic neurons (de Waele et al., 1995; Gliddon et al., 2005), which can themselves be divided in subgroups depending of their morphology, respective networks, or molecular background. When considered in this context, we would expect AAV virus injections into VGluT2-cre and GAD2-cre mice to stimulate only glutamatergic or GABAergic neurons as if each on them belong to a homogeneous neuronal population. This additional layer of complexity cannot be targeted by using this method and will necessitate future investigation to elucidate.

### Functional considerations

It is interesting to note that the posturo-locomotor syndrome induced by unilateral optogenetic stimulation of glutamatergic MVN neurons mirrors that of typical posturo-locomotor behavior evoked by unilateral lesion of the vestibular system with arsanilate. During optogenetic stimulation of MVN glutamatergic neurons, animals rotate towards the side of the simulation, while they circle towards the side ipsilateral to the lesion in the arsanilate model. These typical postural behaviors generated by both optogenetic stimulation and arsanilate lesion result from the electrophysiological asymmetry between ipsi and contralateral MVNs. Indeed, a large number of studies have shown that immediately after unilateral vestibular lesion, resting discharge rates of MVN neurons on the ipsilesional side are abolished, whilst the activity of MVN neurons on the contralesional side is either normal or increased (Newlands and Perachio, 1990; Ris and Godaux, 1998; Smith and Curthoys, 1989). In this situation, the vestibular deficit is always on the side of the body corresponding to the MVN with reduced electrical activity. Muscle hypotension, falls, head tilt and circling affect the side in which the MVN is electrically deficient. Furthermore, the decline of the posturo-locomotor symptoms coincides with a recovery of the resting activity of ipsilesional MVN (Lacour and Tighilet, 2010). The posturo-locomotor deficits consecutive to optogenetic stimulation of glutamatergic MVN neurons likely result from an electrophysiological imbalance with a highly increased activity on the stimulated side.

To better appreciate the behavioral phenotype common to both arsanilate lesion and optogenetic stimulation of VN, it is worth examining VN connectomics. The medial vestibulospinal tract is primarily composed of axons from MVN neurons, but fibers from the lateral and descending vestibular nuclei also contribute to the tract (Carpenter, 1991). Axons of the medial tract travel in the medial longitudinal fasciculus and principally terminate in upper cervical regions of the spinal cord that innervate upper-body musculature, particularly neck musculature. Thus, the MVN and medial vestibulospinal tract provide the circuitry necessary to rapidly control vestibulo-collic reflexes and neck postural changes associated with alterations in head orientation. The lateral vestibulospinal tract is the largest of the two tracts and is primarily composed of axons from neurons in the lateral vestibular nucleus (LVN, also referred as Dieter’s nucleus), with some contribution from the descending (inferior) nucleus. The fibers descend ipsilaterally in the ventrolateral columns and have branches that innervate multiple levels of the spinal cord, thus providing the capacity to modulate spinal motoneuron activity across segments (Abzug et al., 1974). Thus, vestibulospinal pathways are well positioned to modulate the excitability of reflex responses to anticipated or imposed postural displacements. Experiments in animals have shown that electric stimulation of the LVN increases the excitability of extensor motoneurons and decreases those of flexor motoneurons (Grillner et al., 1970; McCall et al., 2017; Wilson and Yoshida, 1969). The facilitatory effect of vestibulospinal inputs on extensor motoneuron excitability suggests that the predominant function of this pathway is to provide appropriate levels of extensor tone to achieve vertical support against gravity. In the present study, the characteristic behaviors observed in optogenetically-stimulated mice, such as tumbling, circling, and retropulsion could be attributed to the specific stimulation of the MVN, whilst the enlargement of the support surface area, and head tilt (observed but not measured) could result from stimulation of the LVN. Most commissural inhibition is provided by type I excitatory neurons unilaterally acting on type II inhibitory neurons on the opposite side, thus releasing GABA in tonic type I neurons on the opposite side (Kasahara et al., 1968; Mano et al., 1968; Precht et al., 1973). The sole parameter that departs from this rule is the circling sense observed after optogenetic stimulation. As recently reported (Marouane et al., 2020), the circling phenotype is common to rodent models of Parkinson’s disease (Smith, 2018). This phenomenon may be explained by the involvement of vestibulo-striatal connections that are essential for the control of postural locomotion (Lopez and Blanke, 2011). Circling results from striatal electrophysiological imbalance common to that induced in the VN by both arsanilate lesion and optogenetic stimulation. Further anatomo-functional study to determine the neurochemical nature of vestibulo-striatal connections should allow determining the bases of circling rotation sense after optogenetic stimulation.

When optogenetic stimulation was interrupted, a restoration of normal postural behavior was observed, likely because electrophysiological balance is restored between opposing MVNs. The advantage of the optogenetic stimulation protocol is that we instantly appreciate the posturo-locomotor recovery phenomenon. In terms of both behavior and neuronal responses, the onset of vestibular compensation occurs later and lasts longer in vestibular lesion models compared to the short recovery times following optogenetic stimulation. The disappearance of behavioral deficits as well as the rebalancing of spontaneous activity in the homologous MVN is observed around one week after unilateral vestibular lesion of the vestibular system in guinea pigs (Ris et al., 1997).

Unlike the glutamatergic neural network, unilateral optogenetic stimulation of GABAergic neurons in the MVN did not produce a posturo-locomotor syndrome. Optogenetic stimulation of the GABAergic neural network should have, in theory, induced ipsilateral inhibition of the MVN, thus generating an ipsilateral postural syndrome similar to that observed after arsanilate injury. One can ask what types of GABAergic MVN neurons were activated by optogenetic stimulation? Answering this question is a challenge considering the complex architectural organization of the MVN GABAergic neural network. Indeed, the GABAergic neurons within the vestibular nuclear complex (VNC) have been classified into five functionally distinct groups (Highstein and Holstein, 2006). Group one consists of commissural GABAergic neurons. Group two consists of GABAergic neurons that project to the oculomotor motoneurons that are involved in the vestibulo-ocular reflex (VOR) (for review see: Beitz and Anderson, 2000). These two groups of neurons are located predominantly in the superior vestibular nucleus (SVN) and the rostral medial vestibular nucleus (MVN) and send projections through the ipsilateral medial longitudinal fasciculus. Group three consists of GABAergic neurons within the MVN, prepositus hypoglossy (PH), and inferior vestibular nucleus (IVN) that project to the nucleus of the inferior olive. Group four consists of GABAergic neurons within the rostral MVN and lateral vestibular nucleus (LVN) that project to the medial vestibulo-spinal tract. Finally, group five consists of local interneurons. To answer the above question, it would be necessary to know which type of GABAergic neurons were the target of optogenetic stimulation. We could potentially refer to a morphological criterion size. Indeed, it has been suggested that neuronal size can indicate differentiate a local GABAergic neuron from a projection neuron (Tighilet et al., 2007). However, neuronal size alone would very likely not be sufficient to select only the type of GABAergic neurons targeted by optogenetic stimulation, since the injected virus would infect all GAD2 neurons and not specific subgroups. It would be useful to investigate the nature of these neurons using neuroanatomical tracing and electrophysiological experiments. Another possible explanation for the absence of a behavioral effect under optogenetic stimulation of GABAergic neurons is the low ratio of GABAergic neurons in the VNC, which is less than 10% of total VNC neurons (Walberg et al., 1990), making it likely that the optogenetic stimulation did not recruit a sufficient number of neurons to generate inhibition of the NVM. Finally, it is possible that optogenetic stimulation targeted vestibulospinal GABAergic neurons with bilateral projections, thus canceling any effect. The vestibular nuclei influence postural control via the medial (mostly from ipsi and contralateral NVM neurons) vestibulospinal tracts (Wilson and Jones, 1979).

Compared with recently documented rat UVN model (Rastoldo et al., 2020), optogenetic stimulation seems to similarly affect parameters such as mean distance moved, velocity and immobility time. In addition, the support surface area is also similarly affected in these three models (Liberge et al., 2010; Marouane et al., 2020). These observations strengthen the optogenetic model as a new vestibular pathological model. Given that efferences arising from the MVePC also project onto oculomotor muscles, in addition to multiple cortical areas, (for review see: Lopez, 2016; Lopez and Blanke, 2011), it is likely that VN unilateral optogenetics stimulation also triggers other functional alterations through modulating the vestibulo-ocular reflex (nystagmus and oscillopsia), vestibulo-sympathetic reflex (increase heart rhythm, drop of body temperature) (Balaban, 1999; Yates et al., 2014), and vestibulo-cortical pathways (spatial disorientation). Further studies are necessary to determine whether these functional consequences reported upon unilateral labyrinthectomy and UVN also take place after optogenetics VN stimulation. The optogenetic approach, which makes it possible to specifically stimulate neurons in different areas of the brain, is suitable for identifying downstream networks originating from the VN, and also the upstream pathways which can regulate function.

### Clinical relevance

The development of an optogenetic approach that selectively stimulates neuronal bundles arising from different zones of the vestibular central pathway is a new step, which will be vital in determining the downstream pathways at the origin of the various symptoms of vestibular syndrome. The transfer of this technology to humans is not yet a reality. However, electrical stimulation of the deep nuclei is already being used clinically for Parkinson’s disease (Gradinaru et al., 2009; Kravitz et al., 2010), and recent advances in the use of optogenetics in humans for therapy raises the possibility of stimulating patients with unilateral or bilateral vestibular areflexia or hyporeflexia, a restoration of the peripheral inputs, through vestibular optic stimulation or implant interface (Guyot and Perez Fornos, 2019; Sluydts et al., 2020). The use of such technological advances is quickly becoming a reality.

## Materials & Methods

### Animals

All procedures were approved by Animal Care and Use Committees in the Shenzhen Institute of Advanced Technology (SIAT), Chinese Academy of Sciences (CAS). Wild type C57BL/6 (6–8 weeks) mice (Beijing Vital River Laboratory Animal Technology Co. Ltd, Beijing, China) were used in this study. In addition, adult (6-8 weeks) male VGluT2-ires-cre (Jax No. 016963) and male GAD2-ires-cre (Jax No. 010802) transgenic mice were used. All mice were maintained on a 12/12-h light/dark cycle at 25 °C. Food and water were available *ad libitum*.

### Viral preparation

For optogenetic experiments, we used one of the following viruses depending on the protocol: AAV9-EF1a-DIO–hChR2(H134R)–mCherry or AAV9-EF1a-DIO–mCherry. Virus titers were approximately 2-6×10^12^ vg/ml. Viruses were purchased from Brain VTA Co., Ltd., Wuhan and Shanghai Taitool Bioscience Co.Ltd.

### Viral injections

VGluT2-ires-cre and GAD2-ires-cre mice were anesthetized with pentobarbital (i.p., 80 mg/kg) and fixed on a stereotaxic apparatus (RWD, Shenzhen, China). During surgeries, mice were kept anesthetized with isoflurane (1%) and placed on a heating pad to maintain a body temperature of 35 °C. A 10-μL micro-syringe with a 33-Ga needle (Neuros; Hamilton, Reno, USA) was connected to a microliter syringe pump (UMP3/Micro4; WPI, USA) and slowly placed for virus injection into the VN (coordinates: AP:–6.0 mm, ML: –1.2 mm and DV:–4.5 mm). Virus was then injected (120 nl at a rate of 50 nl/min). After injection, the needle was kept in the place for an additional 5-10 minutes to facilitate the diffusion of the virus, after which it was slowly withdrawn.

### Implantation of optical fibers

Optical fibers (200 um diameter, NA: 0.37) were chronically implanted in the VN 3 weeks after virus infusion. For optogenetic experiments, an optical fiber was unilaterally implanted above the VN(AP: –6.0 mm, ML: –1.2 mm and DV: –3.8 mm). After surgery all animals were given at least one week to recover before experiments began.

### Histology, immunohistochemistry, and microscopy

Mice were euthanized with an overdose of chloral hydrate (300 mg/kg, i.p) and transcardially perfused with ice-cold 1 × PBS and then with ice-cold 4% paraformaldehyde (PFA, sigma) in 1 x PBS. Brains were extracted and submerged overnight in 4% PFA at 4 °C before being switched to 30% sucrose in 1 × PBS to equilibrate. Brains were cut into 40-μm thick coronal sections using a cryostat microtome (Lecia CM1950, Germany). Finally, slices were mounted and cover-slipped in anti-fade aqueous mounting reagent with DAPI (ProLong Gold Antifade Reagent with DAPI, life technologies). Brain sections were then photographed and analyzed with an Olympus VS120 virtual microscopy slide scanning system and ImageJ and Photoshop software.

### Arsanilate injection

Once mice were deeply anesthetized using pentobarbital (i.p. 80 mg/kg), a micro needle (30 gauge) was used to create a hole in the eardrum and infuse 0.150 ml of a 10% p-arsanilic acid solution (A9258, dissolved in PBS). Following transtympanic instillation, each mouse was laid on its side for 30 minutes with the lesion side was on top.

### Behavior

The behavioral capacities of mice in the arsanilate group was assessed at 7 times points relative to day of surgery: baseline (before surgery), then 24, 48 and 72 hours after, and at 1, 2 and 3 weeks. Animals performing optogenetic stimulations were exposed to the same apparatus as the arsanilate group, and were habituated to the open field one day before the experiment. During the stimulation day, animals were first placed in the field for a 5-minute phase without stimulation, and then optogenetic stimulation (473-nm blue laser; Aurora-220-473, NEWDOON, Hangzhou) was delivered for 30 seconds (VGluT2: 20 Hz, 5 ms, 473 nm; GAD2: 60 Hz, 5 ms, 473 nm), followed by a final 5 minute without stimulation. The locomotion test was performed in an open field (40 cm x 40 cm x 40 cm) to which mice were habituated to the day before the test. During the test, mice were lifted by their tails for 5 quick vertical movements, then were immediately placed in the open field. Behavioral recordings were made by 2 cameras (one top-view, one side-view) for a duration of 1 minute, to assess vestibular symptoms, which include (i) tumbling: spontaneous or evoked rotations of the animals along their body axis, rated 5 in our evaluation scale, (ii) Retropulsion: characterizes backwards movements of the animals, rated 4; (iii) Circling: circular movements of the rats in the horizontal plane, rated 3, (iv) Bobbing: relates to rapid head tilts to the rear and is assimilated to cephalic nystagmus, rated 2, and (v) Head tilt, rated 1. When none of these behaviors were observed, animals received a score of 0.

For the tail-hanging landing paradigm (THL), mice were lifted by their tails for 5 quick vertical movements. This test normally induced forelimb extension as the animals reach the ground. Unilateral vestibular deficit resulted in trouble during the landing process. Such deficits were quoted from 0 (perfect preparation of the two front paws before reaching the ground) to 3 (absence of preparation for landing). The landing process was accompanied by axial rotation of the body, i.e. a twirl movement that we quantified from 0 (no rotation) to 3 (continuous twisting). We also monitored the intensity of syndrome reactivation once landed, from 0 (no sign) to 3 (max expression/accentuation of circling, tumbling, muscle dystonia, bobbing or head tilt). Mice receiving optogenetic stimulation during THL tests were first exposed to the same protocol as arsanilate group (lifted by the tail without additional stimulation). Then they received optogenetic stimulation continuously for 30 s, whilst being tested for lifting and landing. Two cameras recorded,1 from the side,1 from the bottom.

### Data analysis

Speed, immobility, and distance data were analyzed using Anymaze software (Stoelting Co.). Support surface area and video frames were first analyzed using Kinovea software (https://www.kinovea.org/), and then MATLAB was used to compute the size of the surface. Statistical significance was determined using two-tailed Student’s t-tests with a significance level of p<0.05.

## Acknowledgements

This work was supported by Key-Area Research and Development Program of Guangdong Province 2018B030331001; National Natural Science Foundation of China (NSFC) 31930047(to L.W.); NSFC 31630031 (L.W.); NSFC 91732304 (L.W.); the Strategic Priority Research Program of Chinese Academy of Science XDB32030100; Key Laboratory of SIAT 2019DP173024; International Partnership Program of Chinese Academy of Sciences 172644KYS820170004 (L.W.); the CAS President’s International Fellowship for Young Staff 2020FYB0005 (Q.M.); Shenzhen Key Science and Technology Infrastructure Planning Project ZDKJ20190204002; Guangdong Province Basic Research Grant 2019A1515110870 (X.L), 32000711 NSFC (X.L); and PHC XU GUANGQI 2019 program (Project N° 43374UD) from Campus France (Q.M., C.C., B.T., S.B., L.W.).

## Author contributions

**Quentin Montardy:** Conceptualization, methodology, validation, formal analysis, data curation, writing – original draft, Writing - Review & Editing, visualization, project administration **Mengxia Wei:** Investigation (virus injection, fiber implantation, arsanilate injection and behavior, immunohistochemistry and microscopy), data curation and analyzes, writing – original draft, Writing **Tian Yi:** Investigation (virus injection and fiber implantation, arsanilate injection and behavior), data curation and analyzes, writing – original draft **Xuemei Liu:** Investigation (performed immunohistochemistry and quantitative analyzed of the tracing data), manuscript review and editing **Zhou Zheng:** Methodology (setup injection protocol), conceptualization, manuscript review and editing. **Lai Juan:** Investigation (immunohistochemistry) **Stephane Besnard:** Conceptualization, manuscript review and editing **Brahim Tighilet:** Conceptualization, original draft writing, manuscript review and editing **Christian Chabbert:** Conceptualization, supervision, original draft writing, manuscript review and editing **Liping Wang:** Conceptualization, supervision & Project administration, manuscript review and editing

## References

Abzug, C., Maeda, M., Peterson, B.W., Wilson, V.J., Bean, C.P., 1974. Cervical branching of lumbar vestibulospinal axons: With an Appexdix. The Journal of Physiology 243, 499–522. https://doi.org/10.1113/jphysiol.1974.sp010764

Adams, F., 1849. The genuine works of Hippocrates. Sydenham society.

Alvarez, J.C., Díaz, C., Suárez, C., Fernández, J.A., González del Rey, C., Navarro, A., Tolivia, J., 1998. Neuronal loss in human medial vestibular nucleus. Anat. Rec. 251, 431–438. https://doi.org/10.1002/(SICI)1097-0185(199808)251:4<431::AID-AR2>3.0.CO;2-V

Angelaki, D.E., Cullen, K.E., 2008. Vestibular System: The Many Facets of a Multimodal Sense. Annu. Rev. Neurosci. 31, 125–150. https://doi.org/10.1146/annurev.neuro.31.060407.125555

Balaban, C.D., 1999. Vestibular autonomic regulation (including motion sickness and the mechanism of vomiting). Curr. Opin. Neurol. 12, 29–33. https://doi.org/10.1097/00019052-199902000-00005

Balaban, C.D., Beryozkin, G., 1994. Organization of vestibular nucleus projections to the caudal dorsal cap of kooy in rabbits. Neuroscience 62, 1217–1236. https://doi.org/10.1016/0306-4522(94)90354-9

Balaban, C.D., Porter, J.D., 1998. Neuroanatomic substrates for vestibulo-autonomic interactions. J Vestib Res 8, 7–16.

Bárány, R., 1936. Die Beeinflussung des Ohrensausens durch intravenös injizierte Lokalanästhetica. Vorläufige Mitteilung. Acta Oto-Laryngologica 23, 201–203. https://doi.org/10.3109/00016483609123219

Beitz, A.J., Anderson, J.H. (Eds.), 2000. Neurochemistry of the vestibular system. CRC Press, Boca Raton, FL.

Borel, L., Lopez, C., Péruch, P., Lacour, M., 2008. Vestibular syndrome: A change in internal spatial representation. Neurophysiologie Clinique/Clinical Neurophysiology 38, 375–389. https://doi.org/10.1016/j.neucli.2008.09.002

Brugeaud, A., Travo, C., Dememes, D., Lenoir, M., Llorens, J., Puel, J.-L., Chabbert, C., 2007. Control of Hair Cell Excitability by Vestibular Primary Sensory Neurons. Journal of Neuroscience 27, 3503–3511. https://doi.org/10.1523/JNEUROSCI.5185-06.2007

Carpenter, M.B., 1991. Core text of neuroanatomy, 4th ed. ed. Williams & Wilkins, Baltimore.

de Waele, C., Mühlethaler, M., Vidal, P.P., 1995. Neurochemistry of the central vestibular pathways. Brain Research Reviews 20, 24–46. https://doi.org/10.1016/0165-0173(94)00004-9

Desmadryl, G., Gaboyard-Niay, S., Brugeaud, A., Travo, C., Broussy, A., Saleur, A., Dyhrfjeld-Johnsen, J., Wersinger, E., Chabbert, C., 2012. Histamine H _4_ receptor antagonists as potent modulators of mammalian vestibular primary neuron excitability: H _4_ receptor modulates vestibular neuron excitability. British Journal of Pharmacology 167, 905–916. https://doi.org/10.1111/j.1476-5381.2012.02049.x

Dutheil, S., Lacour, M., Tighilet, B., 2011. Neurogenic Potential of the Vestibular Nuclei and Behavioural Recovery Time Course in the Adult Cat Are Governed by the Nature of the Vestibular Damage. PLoS ONE 6, e22262. https://doi.org/10.1371/journal.pone.0022262

Dyhrfjeld-Johnsen, J., Gaboyard-Niay, S., Broussy, A., Saleur, A., Brugeaud, A., Chabbert, C., 2013. Ondansetron reduces lasting vestibular deficits in a model of severe peripheral excitotoxic injury. Journal of Vestibular Research 23, 177–186. https://doi.org/10.3233/VES-130483

Flourens, P., 1842. Recherches expérimentales sur les propriétés et les fonctions du système nerveux dans les animaux vertébrés. Ballière.

Gaboyard-Niay, S., Travo, C., Saleur, A., Broussy, A., Brugeaud, A., Chabbert, C., 2016. Correlation between afferent rearrangements and behavioral deficits after local excitotoxic insult in the mammalian vestibule: a rat model of vertigo symptoms. Dis. Model. Mech. 9, 1181–1192. https://doi.org/10.1242/dmm.024521

Gliddon, C., Darlington, C., Smith, P., 2005. GABAergic systems in the vestibular nucleus and their contribution to vestibular compensation. Progress in Neurobiology 75, 53–81. https://doi.org/10.1016/j.pneurobio.2004.11.001

Gradinaru, V., Mogri, M., Thompson, K.R., Henderson, J.M., Deisseroth, K., 2009. Optical Deconstruction of Parkinsonian Neural Circuitry. Science 324, 354–359. https://doi.org/10.1126/science.1167093

Grillner, S., Hongo, T., Lund, S., 1970. The vestibulospinal tract. Effects on alpha-motoneurones in the lumbosacral spinal cord in the cat. Exp. Brain Res. 10, 94–120. https://doi.org/10.1007/BF00340521

Guyot, J.-P., Perez Fornos, A., 2019. Milestones in the development of a vestibular implant: Current Opinion in Neurology 32, 145–153. https://doi.org/10.1097/WCO.0000000000000639

Hamann, K.F., Reber, A., Hess, B.J.M., Dieringer, N., 1998. Long-term deficits in otolith, canal and optokinetic ocular reflexes of pigmented rats after unilateral vestibular nerve section. Experimental Brain Research 118, 331–340. https://doi.org/10.1007/s002210050287

Highstein, S.M., Holstein, G.R., 2006. The Anatomy of the vestibular nuclei, in: Progress in Brain Research. Elsevier, pp. 157–203. https://doi.org/10.1016/S0079-6123(05)51006-9

Hirvonen, T.P., Minor, L.B., Hullar, T.E., Carey, J.P., 2005. Effects of Intratympanic Gentamicin on Vestibular Afferents and Hair Cells in the Chinchilla. Journal of Neurophysiology 93, 643–655. https://doi.org/10.1152/jn.00160.2004

Hitier, M., Besnard, S., Vignaux, G., Denise, P., Moreau, S., 2010. The ventrolateral surgical approach to labyrinthectomy in rats: anatomical description and clinical consequences. Surg Radiol Anat 32, 835–842. https://doi.org/10.1007/s00276-010-0690-9

Kasahara, M., Mano, N., Oshima, T., Ozawa, S., Shimazu, H., 1968. Contralateral short latency inhibition of central vestibular neurons in the horizontal canal system. Brain Research 8, 376–378. https://doi.org/10.1016/0006-8993(68)90057-7

Kravitz, A.V., Freeze, B.S., Parker, P.R.L., Kay, K., Thwin, M.T., Deisseroth, K., Kreitzer, A.C., 2010. Regulation of parkinsonian motor behaviours by optogenetic control of basal ganglia circuitry. Nature 466, 622–626. https://doi.org/10.1038/nature09159

Lacour, M., Helmchen, C., Vidal, P.-P., 2016. Vestibular compensation: the neuro-otologist’s best friend. J Neurol 263, 54–64. https://doi.org/10.1007/s00415-015-7903-4

Lacour, M., Roll, J.P., Appaix, M., 1976. Modifications and development of spinal reflexes in the alert baboon (Papio papio) following an unilateral vestibular neurotomy. Brain Research 113, 255–269. https://doi.org/10.1016/0006-8993(76)90940-9

Lacour, M., Tighilet, B., 2010. Plastic events in the vestibular nuclei during vestibular compensation: The brain orchestration of a “deafferentation” code. Restorative Neurology and Neuroscience 28, 19–35. https://doi.org/10.3233/RNN-2010-0509

Li, H., Godfrey, D.A., Rubin, A.M., 1995. Comparison of surgeries for removal of primary vestibular inputs: A combined anatomical and behavioral study in rats. Laryngoscope 105, 417–424. https://doi.org/10.1288/00005537-199504000-00015

Liberge, M., Manrique, C., Bernard-Demanze, L., Lacour, M., 2010. Changes in TNFα, NFκB and MnSOD protein in the vestibular nuclei after unilateral vestibular deafferentation. J Neuroinflammation 7, 91. https://doi.org/10.1186/1742-2094-7-91

Lopez, C., 2016. The vestibular system: balancing more than just the body. Current Opinion in Neurology 29, 74–83. https://doi.org/10.1097/WCO.0000000000000286

Lopez, C., Blanke, O., 2011. The thalamocortical vestibular system in animals and humans. Brain Res Rev 67, 119–146. https://doi.org/10.1016/j.brainresrev.2010.12.002

Lopez, C., Falconer, C.J., Deroualle, D., Mast, F.W., 2015. In the presence of others: Self-location, balance control and vestibular processing. Neurophysiologie Clinique/Clinical Neurophysiology 45, 241–254. https://doi.org/10.1016/j.neucli.2015.09.001

Lorente De No, R., 1933. The central projection of the nerve endings of the internal ear. The Laryngoscope.

Mano, N., Oshima, T., Shimazu, H., 1968. Inhibitory commissural fibers interconnecting the bilateral vestibular nuclei. Brain Research 8, 378–382. https://doi.org/10.1016/0006-8993(68)90058-9

Marouane, E., Rastoldo, G., El Mahmoudi, N., Péricat, D., Chabbert, C., Artzner, V., Tighilet, B., 2020. Identification of New Biomarkers of Posturo-Locomotor Instability in a Rodent Model of Vestibular Pathology. Front. Neurol. 11, 470. https://doi.org/10.3389/fneur.2020.00470

McCall, A.A., Miller, D.M., Yates, B.J., 2017. Descending Influences on Vestibulospinal and Vestibulosympathetic Reflexes. Front. Neurol. 8. https://doi.org/10.3389/fneur.2017.00112

Newlands, S.D., Perachio, A.A., 1990. Compensation of horizontal canal related activity in the medial vestibular nucleus following unilateral labyrinth ablation in the decerebrate gerbil: Type I neurons. Exp Brain Res 82, 359–372. https://doi.org/10.1007/BF00231255

O’Neill, A.B., Pan, J.-B., Sullivan, J.P., Brioni, J.D., 1999. Pharmacological evaluation of in vivo model of vestibular dysfunction in the rat. Methods Find Exp Clin Pharmacol 21, 285. https://doi.org/10.1358/mf.1999.21.4.538180

Paterson, J.M., Menzies, J.R.W., Bergquist, F., Dutia, M.B., 2004. Cellular Mechanisms of Vestibular Compensation. Neuroembryol Aging 3, 183–193. https://doi.org/10.1159/000096796

Patk, T., Vassias, I., Vidal, P.P., De Waele, C., 2003. Modulation of the voltage-gated sodium- and calcium-dependent potassium channels in rat vestibular and facial nuclei after unilateral labyrinthectomy and facial nerve transsection: an in situ hybridization study. Neuroscience 117, 265–280. https://doi.org/10.1016/S0306-4522(02)00829-1

Precht, W., Schwindt, P.C., Baker, R., 1973. Removal of vestibular commissural inhibition by antagonists of GABA and glycine. Brain Research 62, 222–226. https://doi.org/10.1016/0006-8993(73)90631-8

Purkinje, J. von, 1819. Beytr/ige zur Kenntniss des Sehens in subjectiver Hinsicht. Prag: Calve.

Rastoldo, G., Marouane, E., El Mahmoudi, N., Péricat, D., Bourdet, A., Timon-David, E., Dumas, O., Chabbert, C., Tighilet, B., 2020. Quantitative Evaluation of a New Posturo-Locomotor Phenotype in a Rodent Model of Acute Unilateral Vestibulopathy. Front. Neurol. 11, 505. https://doi.org/10.3389/fneur.2020.00505

Ris, L., Capron, B., de Waele, C., Vidal, P.P., Godaux, E., 1997. Dissociations between behavioural recovery and restoration of vestibular activity in the unilabyrinthectomized guinea-pig. J. Physiol. (Lond.) 500 (Pt 2), 509–522. https://doi.org/10.1113/jphysiol.1997.sp022037

Ris, L., Godaux, E., 1998. Neuronal Activity in the Vestibular Nuclei After Contralateral or Bilateral Labyrinthectomy in the Alert Guinea Pig. Journal of Neurophysiology 80, 2352–2367. https://doi.org/10.1152/jn.1998.80.5.2352

Sluydts, M., Curthoys, I., Vanspauwen, R., Papsin, B.C., Cushing, S.L., Ramos, A., Ramos de Miguel, A., Borkoski Barreiro, S., Barbara, M., Manrique, M., Zarowski, A., 2020. Electrical Vestibular Stimulation in Humans: A Narrative Review. Audiol Neurotol 25, 6–24. https://doi.org/10.1159/000502407

Smith, P.F., 2018. Vestibular Functions and Parkinson’s Disease. Front. Neurol. 9, 1085. https://doi.org/10.3389/fneur.2018.01085

Smith, P.F., Curthoys, I.S., 1989. Mechanisms of recovery following unilateral labyrinthectomy: a review. Brain Research Reviews 14, 155–180. https://doi.org/10.1016/0165-0173(89)90013-1

Straka, H., Vibert, N., Vidal, P.P., Moore, L.E., Dutia, M.B., 2005. Intrinsic membrane properties of vertebrate vestibular neurons: Function, development and plasticity. Progress in Neurobiology 76, 349–392. https://doi.org/10.1016/j.pneurobio.2005.10.002

Tighilet, B., Brezun, J.M., Dit Duflo Sylvie, G., Gaubert, C., Lacour, M., 2007. New neurons in the vestibular nuclei complex after unilateral vestibular neurectomy in the adult cat: Reactive neurogenesis in adult vestibular lesioned cats. European Journal of Neuroscience 25, 47–58. https://doi.org/10.1111/j.1460-9568.2006.05267.x

Vignaux, G., Chabbert, C., Gaboyard-Niay, S., Travo, C., Machado, M.L., Denise, P., Comoz, F., Hitier, M., Landemore, G., Philoxène, B., Besnard, S., 2012. Evaluation of the chemical model of vestibular lesions induced by arsanilate in rats. Toxicology and Applied Pharmacology 258, 61–71. https://doi.org/10.1016/j.taap.2011.10.008

Walberg, F., Ottersen, O.P., Rinvik, E., 1990. GABA, glycine, aspartate, glutamate and taurine in the vestibular nuclei: an immunocytochemical investigation in the cat. Exp Brain Res 79. https://doi.org/10.1007/BF00229324

Wilson, V.J., Jones, G.M., 1979. Mammalian Vestibular Physiology. Springer US: Imprint: Springer, Boston, MA.

Wilson, V.J., Yoshida, M., 1969. Comparison of effects of stimulation of Deiters’ nucleus and medial longitudinal fasciculus on neck, forelimb, and hindlimb motoneurons. Journal of Neurophysiology 32, 743–758. https://doi.org/10.1152/jn.1969.32.5.743

Xerri, C., Lacour, M., 1980. [Compensation deficits in posture and kinetics following unilateral vestibular neurectomy in cats. The role of sensorimotor activity]. Acta Otolaryngol. 90, 414–424.

Xia, L., Chen, Z., Yin, S., 2012. Ototoxicity of cisplatin administered to guinea pigs via the round window membrane. J Toxicol Sci 37, 823–830.

Yates, B.J., Bolton, P.S., Macefield, V.G., 2014. Vestibulo-Sympathetic Responses, in: Terjung, R. (Ed.), Comprehensive Physiology. John Wiley & Sons, Inc., Hoboken, NJ, USA, pp. 851–887. https://doi.org/10.1002/cphy.c130041

Zhang, M., Liu, W., Ding, D., Salvi, R., 2003. Pifithrin-α supresses p53 and protects cochlear and vestibular hair cells from cisplatin-induced apoptosis. Neuroscience 120, 191–205. https://doi.org/10.1016/S0306-4522(03)00286-0

